# Scallop2 enables accurate assembly of multiple-end RNA-seq data

**DOI:** 10.1101/2021.09.03.458862

**Authors:** Qimin Zhang, Qian Shi, Mingfu Shao

## Abstract

Transcript assembly (i.e., to reconstruct the full-length expressed transcripts from RNA-seq data) has been a critical but yet unsolved step in RNA-seq analysis. Modern RNA-seq protocols can produce paired-/multiple-end RNA-seq reads, where information is available that two or more reads originate from the same transcript. The long-range constraints implied in these paired-/multiple-end reads can be much beneficial in correctly phasing the complicated spliced isoforms. However, there often exist gaps among individual ends, which may even contain junctions, making the efficient use of such constraints algorithmically challenging. Here we introduce Scallop2, a new reference-based transcript assembler optimized for multiple-end (including paired-end) RNA-seq data. Scallop2 uses an algorithmic framework that first represents reads from the same molecule as the so-called multiple-end phasing paths in the context of a splice graph, then “bridges” each multiple-end phasing path into a long, single-end phasing path, and finally decomposes the splice graph into paths (i.e., transcripts) guided by the bridged phasing paths. An efficient bridging algorithm is designed to infer the true path connecting two consecutive ends following a novel formulation that is robust to sequencing errors and transcript noises. By observing that failing to bridge two ends is mainly due to incomplete splice graphs, we propose a new method to determine false starting/ending vertices of the splice graphs which has been showed efficient in reducing false positive rate. Evaluations on both (multiple-end) single-cell RNA-seq datasets from Smart-seq3 protocol and Illumina paired-end RNA-seq samples demonstrate that Scallop2 vastly outperforms recent assemblers including StringTie2, Scallop, and CLASS2 in assembly accuracy.

## 1 Introduction

The established high-throughput RNA sequencing technologies (RNA-seq) have fueled many biological and biomedical discoveries. One of the critical steps and yet the major computational challenges in RNA-seq analysis is *transcript assembly*—the reconstruction of full-length expressed transcripts captured by a given RNA-seq sample. Transcript assembly is the main approach to determine novel transcripts and to annotate gene structures. It also enables the construction of transcriptomes at species-, tissue-, and cell-specific level, by leveraging large-scale available RNA-seq data [1]. Such data-driven transcriptomic infrastructures are crucial for downstream expression quantification and differential analysis.

Over the decades, tremendous efforts have been made to develop efficient approaches for assembling various types of RNA-seq data. For short-reads RNA-seq data (typically Illumina RNA-seq data), Cufflinks [2], Scripture [3], Traph [4], CLASS2 [5], TransComb [6], Scallop [7], StringTie [8], and RefShannon [9] have been developed. For assembling long-reads RNA-seq data (e.g., generated by PacBio Iso-Seq and Oxford Nanopore direct RNA and cDNA sequencing), StringTie2 [10] and Scallop-LR [11] have been released. The recently developed protocols such as Smart-seq series [12, 13] can produce single-cell RNA-seq data with coverage spanning full-length molecules (in contrast to protocols that captures 5’ end of mRNAs), which therefore can be used to assemble transcripts at single-cell resolution. Existing assemblers for such data include scRNAss [14] and RNA-Bloom [15]. We note that the above mentioned tools do not form an exhaustive list and one assembler might be applicable to multiple types of RNA-seq data.

Despite of these, the task of transcript assembly is far from being solved. Assemblers for short-reads RNA-seq data have been struggling with correctly phasing the complete intron-chains from (short) reads especially when genes expressing numerous alternatively spliced isoforms. It remains algorithmically challenging to efficiently use the paired-end information to resolve such complicated cases. The coverage information is helpful in improving phasing, but it is often noisy and biased. Long-reads RNA-seq data has its own limitation of low coverage, making it less efficient in identifying lowly expressed transcripts. A large portion of long reads remain transcript fragments due to cDNA synthesis and sequencing length limit [16], and hence the assembly of long-reads data faces the same challenge as short-reads data. According to benchmarking studies [17] and the performance reported in recent assemblers, the accuracy of existing methods remains unsatisfactory; improved computational methods are therefore still needed for transcript assembly.

Barcoding technologies have been recently incorporated into various sequencing protocols, where the same barcode are appended to the sequencing reads originated from the same molecule or the same fragment of a molecule. Representatives include Linked-Reads protocols developed by 10x Genomics [18, 19] and Smart-seq3 protocol [13]. We refer to the sequencing data generated by these protocols as *multiple-end* reads. The fact that multiple reads are from the same fragment/molecule provides useful long-range information for scaffolding (in genome assembly) and for phasing the intron chains of expressed isoforms (in transcript assembly). Assemblers such as ARBitR [20] and Thena [21] have been developed for multiple-end genomic data. The assembly of multiple-end RNA-seq data is more challenging as there might be multiple ways of filling the gaps between consecutive ends due to the existence of alternative splicing. Efficient algorithms are lacking for assembling multiple-end RNA-seq data.

We present Scallop2, a reference-based transcript assembler for multiple-end RNA-seq data. To make use of the long-range constraints in the data, Scallop2 proposes a new algorithm that infers a single (long) path in the underlying splice graph that connects all individual ends in a read group (i.e., reads with the same barcode). It then employs a “phase-preserving” graph-decomposition algorithm that originally developed in Scallop [7] to decompose the splice graph into paths (i.e., transcripts) while all the inferred long paths are fully preserved. This algorithmic framework is general for assembling data with arbitrary number of ends in a group and is therefore applicable to paired-end RNA-seq data (where each group contains two reads).

## 2 Results

### Resulting tool: Scallop2

We have implemented the algorithm described in Section 3 as a new transcript assembler named Scallop2, freely available at https://github.com/Shao-Group/Scallop2 and built as a conda package at https://anaconda.org/bioconda/scallop2. The input for Scallop2 is the alignments of RNA-seq reads, in standard sam/bam format; the output is the full-length expressed transcripts, in standard gtf format. (If -f option is provided, Scallop2 will also generate the transcript fragments; see Section 3.5.) Scripts that reproduces the experimental results of this paper is available at https://github.com/Shao-Group/scallop2-test.

### Methods compared

We compare Scallop2 (version 1.1.1) with three other recent reference-based transcript assemblers, StringTie2 (version 2.1.7), Scallop (version 0.10.5), and CLASS2 (version 2.1.7). All four assemblers were run with their default parameters. We tried to include more assemblers in the comparison including scRNAss and RNA-Bloom but failed; see Appendix for more details about the efforts we made.

### Datasets

Above four methods are compared with two types of RNA-seq data, generated by Smart-seq3 protocol and Illumina platforms. Smart-seq3 is a single-cell protocol that generates multiple-end RNA-seq data using barcoding technology. Here we use two public datasets published with Smart-seq3 paper [13], downloaded from https://www.ebi.ac.uk/arrayexpress/experiments/E-MTAB-8735. The first dataset, referred to as HEK293T, contains 192 human cells; the second dataset, referred to as Mouse-Fibroblast, includes 369 mouse cells. For Illumina platform, we use 10 human paired-end RNA-seq samples that were downloaded from ENCODE project and were previously used in the Scallop paper [7]. We refer to these 10 samples as ENCODE10. The alignments of these samples are available at http://doi.org/10.26208/8c06-w247.

### Pipeline and evaluation

We use the pipeline illustrated in Figure 1 to evaluate the performance of all compared assemblers. For Smart-seq3 data, the raw sequencing data will be demultiplexed and preprocessed using zUMIs tool, in which STAR will be called to generate reads alignments. The 10 paired-end Illumina RNA-seq samples are aligned with two popular aligners, STAR [22] and HISAT2 [23]. The read alignments of each individual cell or sample will be piped to the compared assemblers.

**Figure 1:**
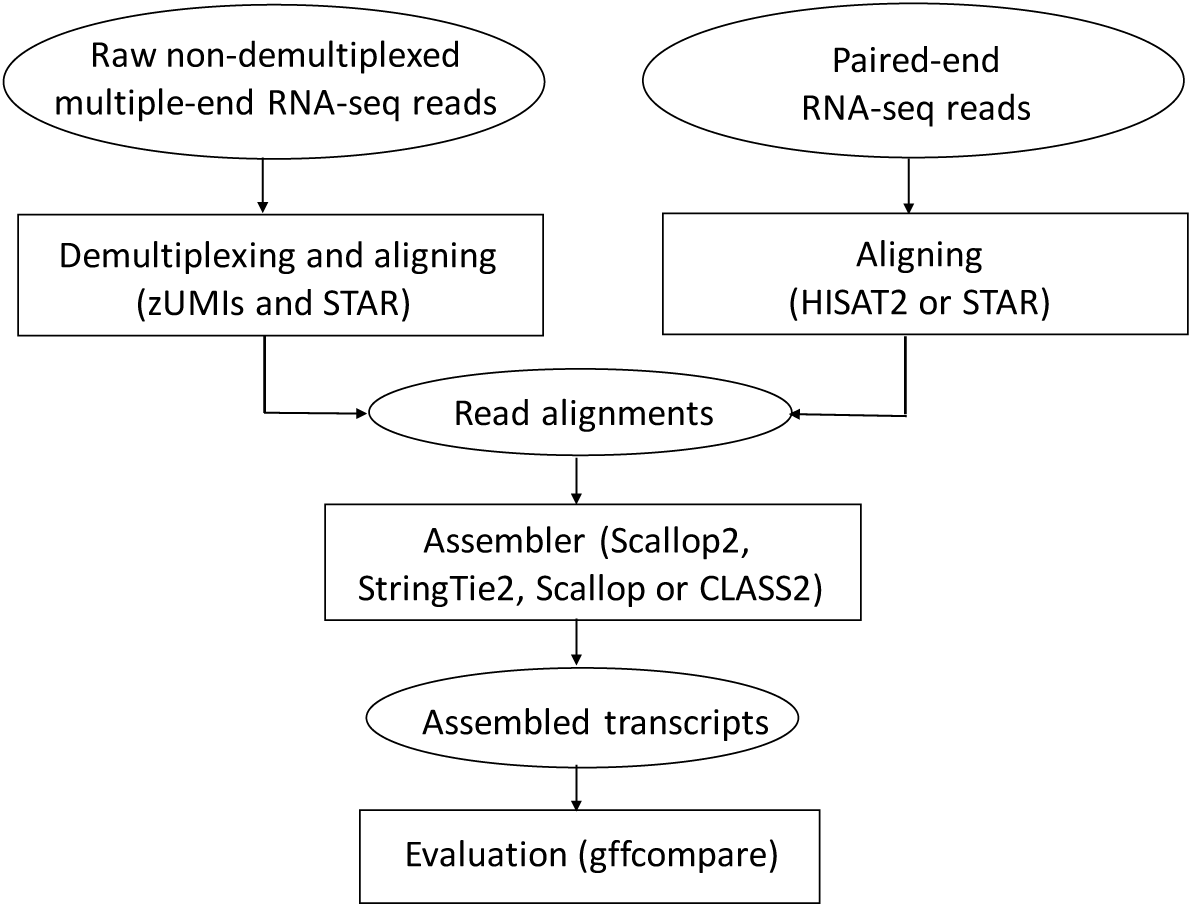
Pipeline of evaluating the performance of compared assemblers.

The accuracy of the assembled transcripts will be evaluated using gffcompare [24]. We use the annotated transcriptomes as reference (Ensembl GRCh38.104 for human data and Ensembl GRCm38.104 for mouse data). We use the “transcript level” metrics defined by gffcompare: an assembled multiple-exon transcript is considered as “matching” if its intron-chain exactly matches that of a transcript in the reference; an assembled single-exon transcript is defined as “matching” if there is a significant overlap (80% by default) with a single-exon transcript in the reference. We use two metrics calculated by gffcompare: the total number of matching transcripts, which is proportional to sensitivity, and precision, defined as the ratio between the total number of matching transcripts and the total number of assembled (predicted) transcripts.

We note that above metrics measured w.r.t. the current transcriptome underestimates the accuracy, as novel transcripts that are correctly assembled will be considered as “not-matching” because they do not yet exist in the reference. Nevertheless, we believe such metrics are fair for the compared assemblers, as they reflect their *relative* accuracy, and yet we do not know the true expressed transcripts in these biological samples.

### Comparison of assembly accuracy on Smart-seq3 data

Table 1 summarizes the assembly accuracy of the 4 assemblers averaged over all cells in each of the two Smart-seq3 datasets. Scallop2 consistently achieved the highest precision and sensitivity on both datasets. Specifically, Scallop2 produces 0.7%, 29.0%, and 20.7% more matching transcripts than StringTie2, Scallop, and CLASS2 on the HEK293T dataset, and assembles 0.4%, 18.1%, and 19.1% more matching transcripts than them on the Mouse-Fibroblast dataset. In terms of precision, Scallop2 improves 107.1%, 12.5% and 119.7% than StringTie2, Scallop, and CLASS2 on the HEK293T dataset, and improves 80.0%, 6.7%, and 66.6% than them on the Mouse-Fibroblast dataset.

**Table 1:**
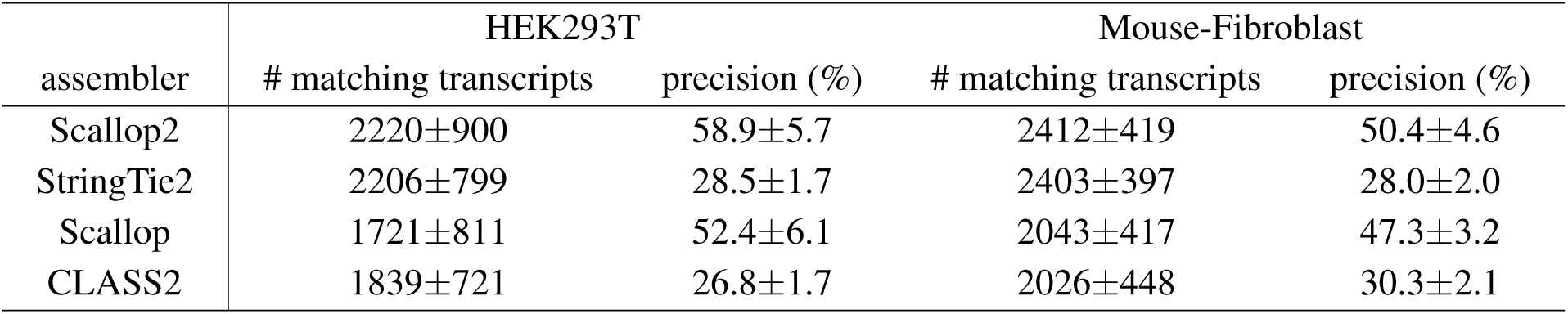
Assembly accuracy of the four assemblers on Smart-seq3 data. The mean and standard deviation are reported over all cells in each dataset.

| assembler | HEK293T |  | Mouse-Fibroblast |  |
| --- | --- | --- | --- | --- |
|  | # matching transcripts | precision (%) | # matching transcripts | precision (%) |
| Scallop2 | 2220±900 | 58.9±5.7 | 2412±419 | 50.4±4.6 |
| StringTie2 | 2206±799 | 28.5±1.7 | 2403±397 | 28.0±2.0 |
| Scallop | 1721±811 | 52.4±6.1 | 2043±417 | 47.3±3.2 |
| CLASS2 | 1839±721 | 26.8±1.7 | 2026±448 | 30.3±2.1 |

Figures 2 and 3 give a direct comparison at the level of individual cells. Scallop2 achieves the highest precision on almost all cells (left panels of Figures 2 and 3), followed by Scallop but vastly outperforms StringTie2 and CLASS2. In terms of sensitivity (right panels of Figures 2 and 3), Scallop2 and StringTie2 significantly outperform the other two methods: in general Scallop2 gives higher sensitivity on cells where all methods exhibits high sensitivity, while StringTie2 wins on cells where methods give low sensitivity. Note though, on average over all cells, Scallop2 slightly outperforms StringTie2 in sensitivity (Table 1).

**Figure 2:**
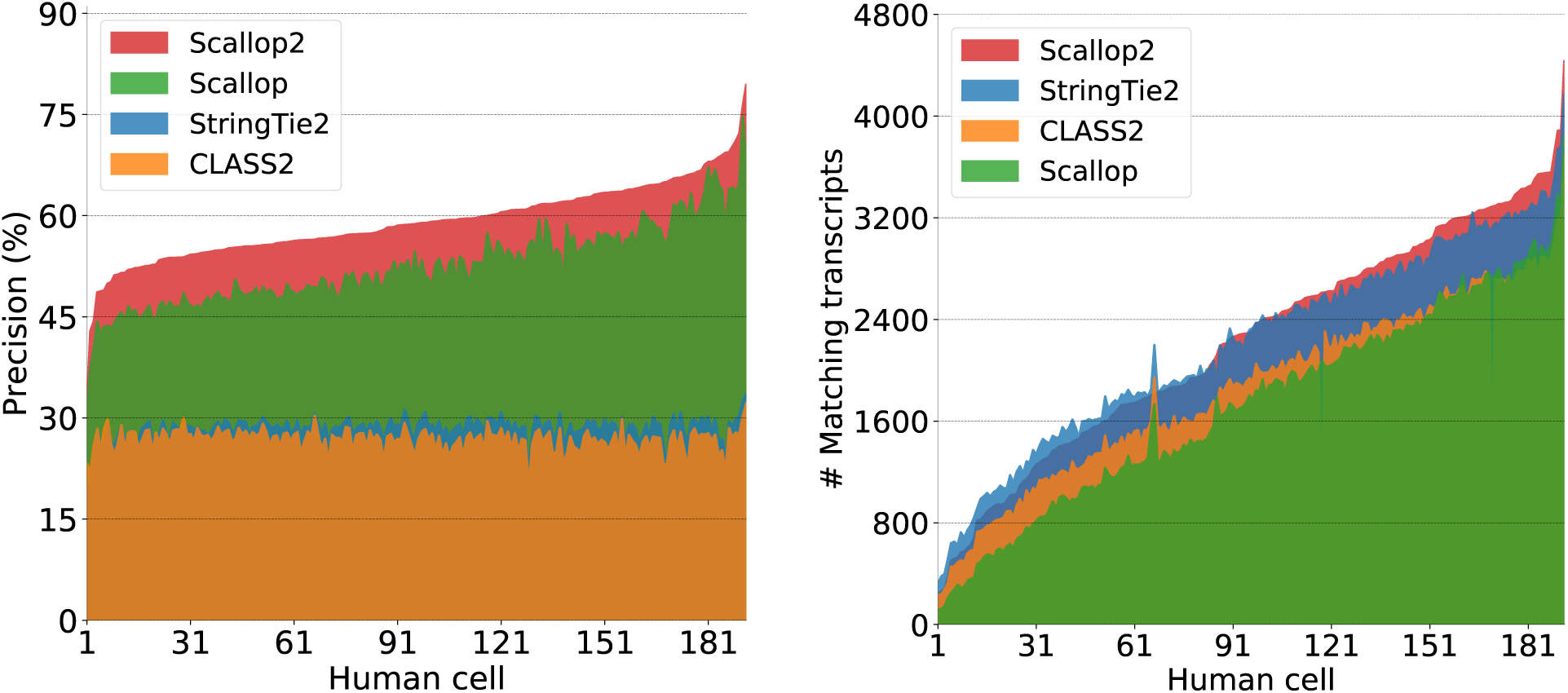
Assembly accuracy at individual cells of HEK293T dataset. In each panel, cells are sorted in ascending order w.r.t. the accuracy of Scallop2 (so the curves for Scallop2 are monotonic).

**Figure 3:**
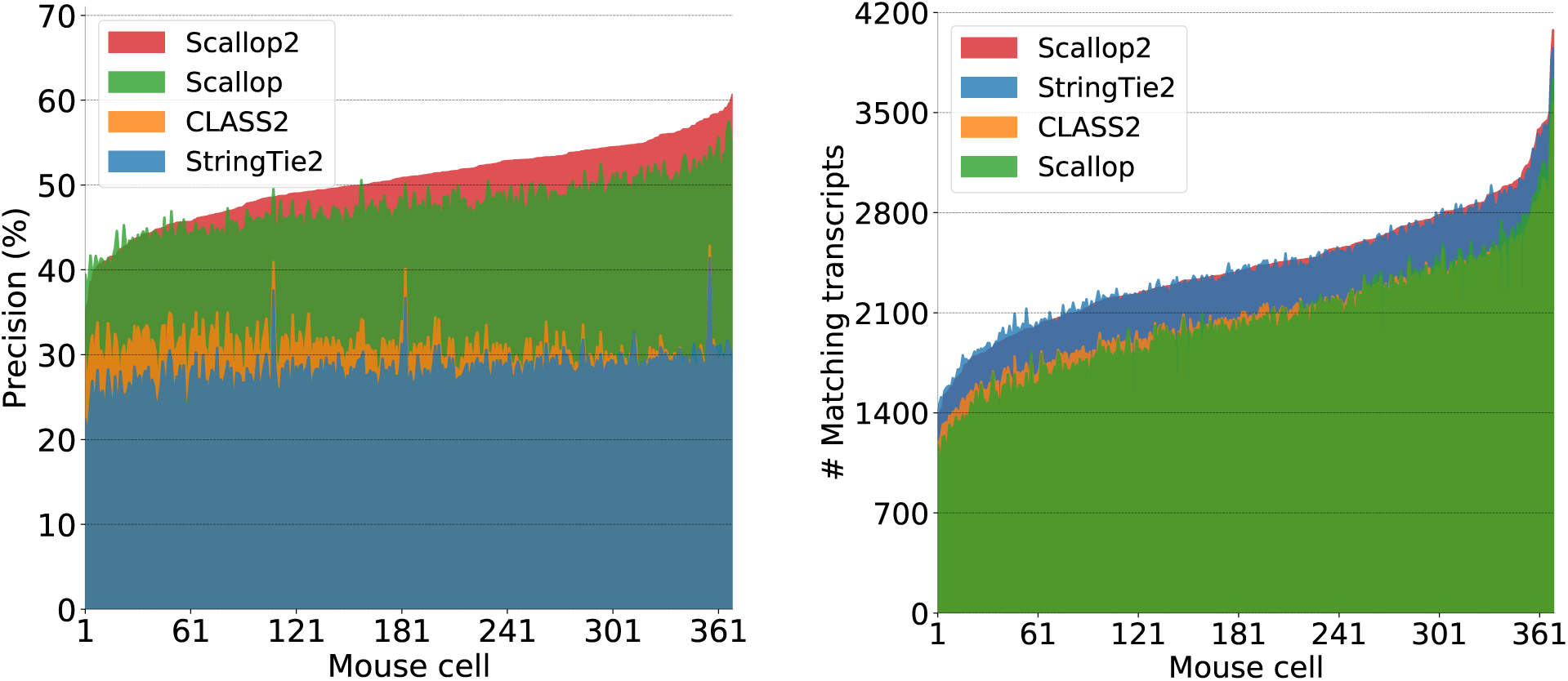
Assembly accuracy at individual cells of Mouse-Fibroblast dataset. In each panel, cells are sorted in ascending order w.r.t. the accuracy of Scallop2 (so the curves for Scallop2 are monotonic).

### Comparison of assembly accuracy on Illumina paired-end RNA-seq data

Table 2 summarizes the assembly accuracy of the four methods on ENCODE10 dataset aligned by HISAT2 and STAR. On average Scallop2 achieved the highest precision and the highest sensitivity with both aligners. Concretely, Scallop2 produces 22.0%, 8.3% and 41.6% more matching transcripts than StringTie2, Scallop and CLASS2 with HISAT2, and produces 23.9%, 5.4% and 38.8% more matching transcripts than them with STAR. In terms of precision, Scallop2 improves 2.7%, 5.7% and 22.8% than StringTie2, Scallop and CLASS2 with HISAT2, and improves 5.8%, 6.1% and 17.6% than them with STAR.

**Table 2:** Assembly accuracy of the four assemblers on ENCODE10 dataset. For each aligner used the mean and standard deviation are reported over the 10 samples.

| assembler | HISAT2 alignments |  | STAR alignments |  |
| --- | --- | --- | --- | --- |
|  | # matching transcripts | precision (%) | # matching transcripts | precision (%) |
| Scallop2 | 17461±1949 | 32.3±7.7 | 17418±2088 | 33.8±8.1 |
| StringTie2 | 14317±2027 | 31.5±6.7 | 14061±2080 | 31.9±6.7 |
| Scallop | 16116±1896 | 30.6±8.4 | 16519±2226 | 31.8±6.9 |
| CLASS2 | 12328±1655 | 26.3±7.8 | 12553±1942 | 28.7±9.1 |

Figures 4 compares the assembly accuracy at individual samples. Scallop2 achieved the highest sensitivity on all samples with both aligners, and obtained the highest precision on 5 (out of 10) samples with HISAT2 and on 6 (out of 10) samples with STAR. To compare on samples where Scallop2 gives higher sensitivity but lower precision, we draw the precision-sensitivity curve of Scallop2. The curve is obtained by gradually filtering out the assembled transcripts with the lowest abundance (i.e., *f* (*p*), predicted by Scallop2 itself); as such filtering goes, the number of matching transcripts of Scallop2 will drop while its precision will likely increase (as lowly-expressed ones are more likely to be false transcripts). Figure 4 shows that, the precision-recall curves of Scallop2 always lie to the right of other points, suggesting that Scallop2 always gives higher precision at the same level of sensitivity compared with any other method.

**Figure 4:**
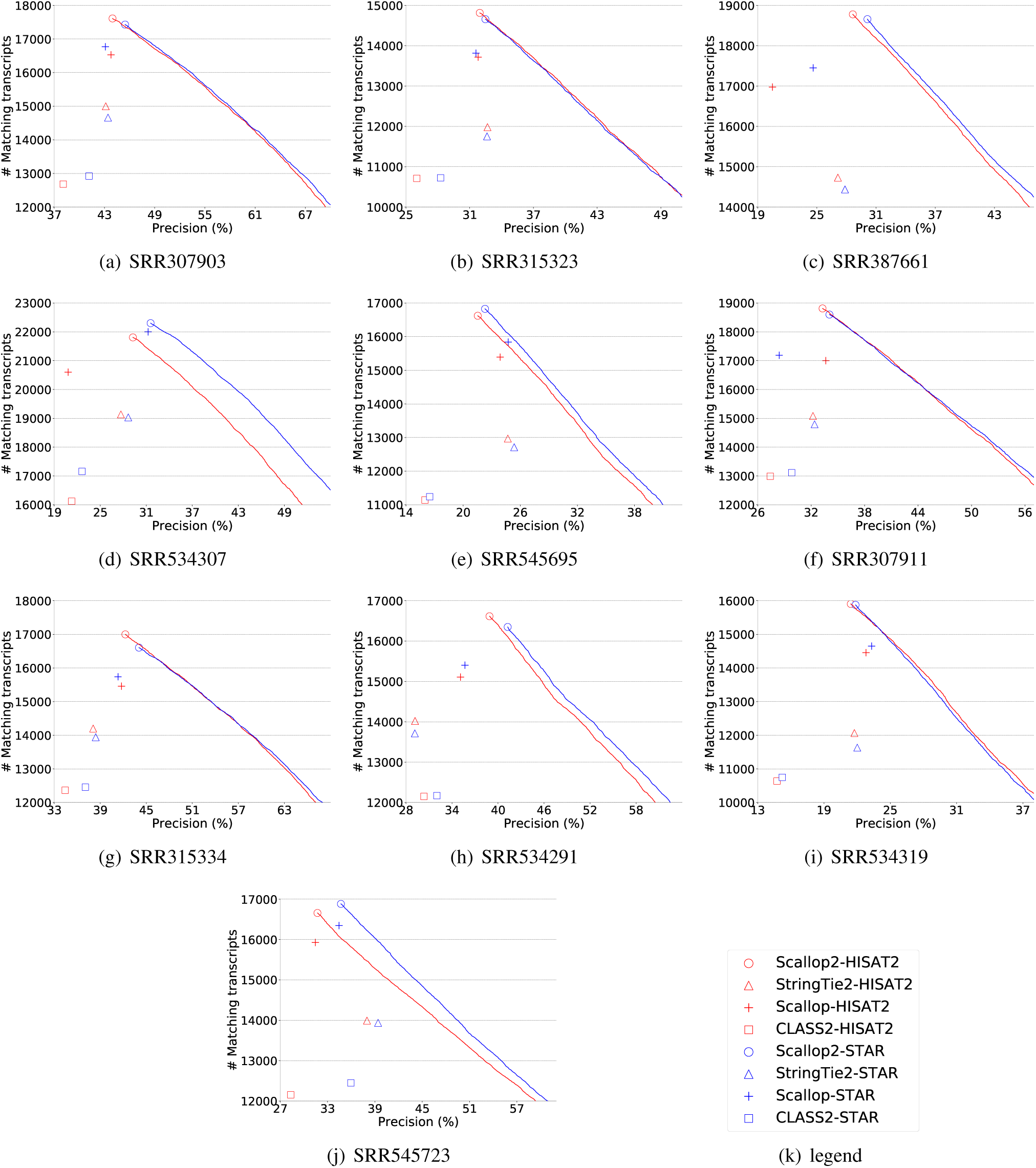
Assembly accuracy on ENCODE10 dataset. CLASS2 did not finish running in 11 days on sample SRR387661.

To quantitatively measure of the improvement of Scallop2 compared with other methods, we calculate the *adjusted precision* of Scallop2 w.r.t. any other method, defined as the *x*-coordinate of the point on the precision-sensitivity curve that has the same *y*-coordinate with the compared method. This single metric therefore offers a more direct evaluation between two methods (as opposed to using two metrics precision and sensitivity). Table 3 summarizes such comparison. When adjusted to the same level of sensitivity, in precision Scallop2 improves 44.6%, 25.1%, and 104.1% than StringTie2, Scallop, and CLASS2 with HISAT2, and improves 49.4%, 18.2%, and 88.2% than them with STAR.

**Table 3:** Comparison of precision (mean and standard deviation) between Scallop2 and each of other 3 methods at the same level of sensitivity. “compared” = (raw) precision of the compared method; “adjusted” = adjusted precision of Scallop2 at the same level of sensitivity with the compared assembler.

| assembler | HISAT2 alignments |  | STAR alignments |  |
| --- | --- | --- | --- | --- |
|  | compared (%) | adjusted (%) | compared (%) | adjusted (%) |
| StringTie2 | 31.5±6.7 | 45.5±8.7 | 31.9±6.7 | 47.7±8.8 |
| Scallop | 30.6±8.4 | 38.2±8.7 | 31.8±6.9 | 37.6±8.5 |
| CLASS2 | 26.3±7.8 | 53.7±10.7 | 28.7±9.1 | 54.1±10.8 |

### Comparison of running time and memory footprint

The comparison of CPU time and peak memory is reported in Table 4 and Table 5. StringTie runs the fastest on Illumina data and Scallop leads on Smart-seq3 data; CLASS2 runs the slowest on both types of data. In terms of memory footprint, StringTie2 significantly outperforms other methods in all cases.

**Table 4:** Comparison of CPU time and peak memory of the 4 methods on Smart-seq3 data. For each dataset, mean and standard deviation over all cells are reported.

| assembler | HEK293T |  | Mouse-Fibroblast |  |
| --- | --- | --- | --- | --- |
|  | running time (s) | memory (MB) | running time (s) | memory (MB) |
| Scallop2 | 15.1±8.9 | 49.0±25.8 | 22.4±9.3 | 73.1±30.6 |
| StringTie2 | 8.0±3.7 | 15.5±4.9 | 10.3±2.8 | 29.5±10.3 |
| Scallop | 4.2±2.4 | 45.5±25.3 | 6.3±2.6 | 62.9±28.1 |
| CLASS2 | 141.2±51.3 | 545.8±19.4 | 102.3±22.9 | 543.7±8.6 |

**Table 5:** Comparison of CPU time and peak memory of the 4 methods on the ENCODE10 dataset. For each aligner used, mean and standard deviation over the 10 samples are reported.

| assembler | HISAT2 alignments |  | STAR alignments |  |
| --- | --- | --- | --- | --- |
|  | running time (s) | memory (MB) | running time (s) | memory (MB) |
| Scallop2 | 1878±1539 | 6124±5072 | 1593±1322 | 6043±4774 |
| StringTie2 | 482±195 | 148±64 | 475±208 | 250±112 |
| Scallop | 949±937 | 8475±9426 | 776±545 | 7839±7960 |
| CLASS2 | 360869±433733 | 2429±1108 | 8596±7305 | 2045±478 |

Scallop2 runs reasonably fast and uses acceptable memory; for example, on average over the 10 Illumina RNA-seq samples aligned with HISAT2, Scallop2 takes about 30 minutes (while the fastest StringTie2 takes about 8 minutes) and uses about 6GB memory (while StringTie2 uses 148MB).

## 3 Methods

We focus on reference-based transcript assembly. The RNA-seq reads will be first aligned to the reference genome through a splice-aware aligners such as STAR [22] and HISAT2 [23]. Reads that overlap in their aligned genomic coordinates are then partitioned as *gene loci*. Individual gene locus will be assembled separately as independent instances. In Section 3.1, we describe the procedure we use to construct the data structures that capture all splicing signals and multiple-end constraints in the data. In Section 3.2, we formulate the multiple-end assembly problem. In Section 3.3, we describe our algorithmic framework for solving above formulation. Two core algorithms called in this framework are described in Section 3.4 (i.e., to bridge two consecutive ends in a group) and in Section 3.5 (i.e., to determine false starting/ending vertices in the splice graph).

### 3.1 Construction of the weighted splice graph and associated multiple-end phasing paths

For each gene locus, we build a weighted splice graph *G* = (*V, E, w*) and a set of multiple-end phasing paths *𝒞* as follows. See Figure 5. Splicing positions are first extracted from the junctions (marked with ‘N’ in the CIGAR string of alignments) to obtain the boundaries of exons (or partial exons) and introns of the reference genome. For each inferred exon, we create a vertex *v* to vertex-set *V*. A directed edge *e* = (*u, v*) is added to edge-set *E* when there exists reads connecting exons *u* and *v*, where *u* occurs before *v* in the genome. The weight of edge *e*, denoted as *w*(*e*), is calculated as the number of reads that connect *u* and *v*.

**Figure 5:**
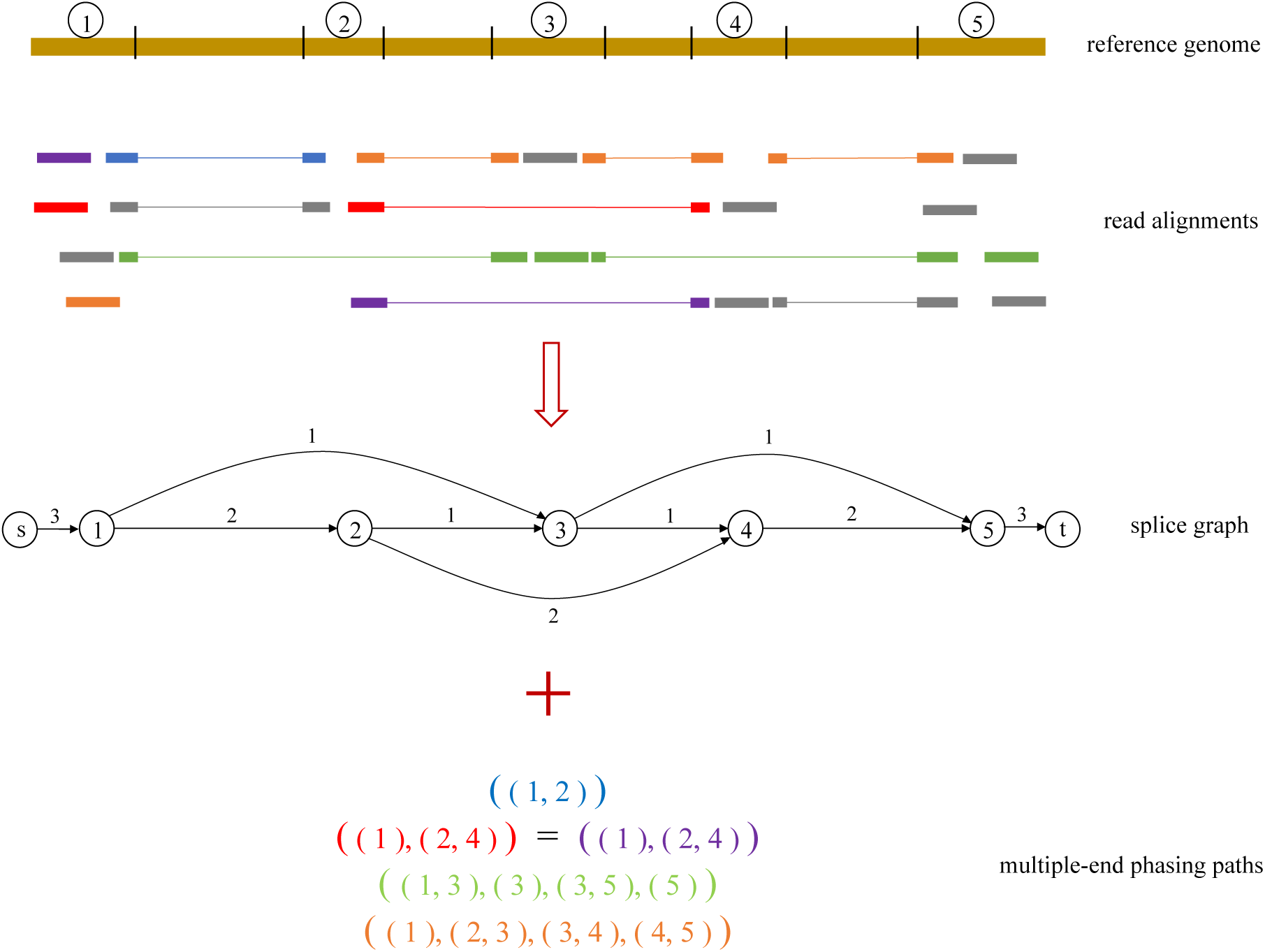
Illustrating the construction of the splice graph *G* and the associated multiple-end phasing paths *𝒞* from read alignments of a gene locus. Reads with the same color represent they form a group (i.e., attached with the same barcode or being the two ends of a pair in paired-end reads); we use gray color to represent reads forming groups of their own.

Additionally, we add a source vertex *s* and sink vertex *t* to *V* ; for each vertex *u* ∈ *V* \{*s,t*} with in-degree of 0 (called *starting* vertices), we add a directed edge (*s, u*) with weight *w*(*s, u*) = ∑_(*u,v*)∈*E*_ *w*(*u, v*), and for each vertex *v* ∈ *V* \{*s,t*} with out-degree of 0 (called *ending* vertices), we add a directed edge (*v,t*) with weight *w*(*v,t*) = ∑_(*u,v*)∈*E*_ *w*(*u, v*).

With the weighted splice graph *G* = (*V, E, w*) being constructed, each read *r* can be represented as a path in *G*, denoted as *l*(*r*), which is the list of vertices of *G* where *r* is aligned to. Let *R* be a set of reads from the same transcript/fragment; we call reads in *R* form a *read group*. In case of barcoding-based prototols like Smart-seq3, reads in group *R* include those attached with the same barcode; in case of paired-end RNA-seq prototols, the two ends of a pair form a group (i.e., |*R*| = 2); in case a read is not attached with any barcode, or its mate end is not aligned, then this read forms a group of its own. Notice that different reads in *R* might be aligned to the same path in *G*, i.e., *l*(*r*_1_) = *l*(*r*_2_) for two reads *r*_1_, *r*_2_ ∈ *R*. We define the *multiple-end phasing path* obtained from *R*, denoted as *C*(*R*), as the set of distinct paths collected from reads in *R*. Formally, *C*(*R*) := {*l*(*r*) | *r* ∈ *R*}. Let {*R*_1_, *R*_2_, …, *R*_*k*_} be the collection of read groups in a gene locus. Again different groups of reads may correspond to the same multiple-end phasing path, i.e., *C*(*R*_*i*_) = *C*(*R*_*j*_) for some 1 ≤ *i < j* ≤ *k*. We denote by *𝒞* as the collection of distinct multiple-end phasing paths in {*C*(*R*_1_),*C*(*R*_2_), …, *C*(*R*_*k*_)}.

### 3.2 Problem formulation

For a gene locus, the above constructed weighted splice graph *G* together with the multiple-end phasing paths *𝒞* give a complete representation of the splicing information in the alignments and the long-range phasing information in the multiple-end reads. Our assembly algorithm will take *G* and *𝒞* as input, and produces a set of *s*-*t* paths *P* of *G* and assigns an abundance *f* (*p*) for each *s*-*t* path *p* ∈ *P*: each *s*-*t* path *p* infers an expressed transcript in this gene locus, and *f* (*p*) predicts its expression abundance.

We now design three objectives to guide reconstructing *P* and *f* given *G* and *𝒞*. First, since each multiple-end phasing path is constructed from reads sampled from a single transcript, we expect that each multiple-end phasing path appears in one of the reconstructed transcripts (i.e., in *P*). Formally, we say a multiple-end phasing path *C* ∈ *𝒞* is *covered* by *P*, if there exists an *s*-*t* path *p* ∈ *P* such that every path *l* ∈ *C* appears in *p*. We require all multiple-end phasing paths in *𝒞* be covered by *P*. Second, for each edge *e* ∈ *E*, we expect the abundance of the inferred *s*-*t* paths passing through *e*, i.e., ∑_*p*∈*P*:*e*∈*p*_ *f* (*p*), is as close to its observed read coverage *w*(*e*) as possible, which can be define as to minimize the sum of the deviation between *w*(*e*) and ∑_*p*∈*P*:*e*∈*p*_ *f* (*p*) for all edges, denoted as *d*(*P, f*) := ∑_*e*∈*E*_ |*w*(*e*)−∑_*p*∈*P*:*e*∈*p*_ *f* (*p*)|. Third, we aim to minimize |*P*| following the parsimony principle. Combining all the three objectives, we informally describe the task of transcript assembly for multiple-end RNA-seq data as follows.

#### Problem 1 (multiple-end transcript assembly)

*Given weighted splice graph G* = (*V, E, w*) *and associated multiple-end phasing paths 𝒞, to compute a set of s-t paths P of G and abundance f* (*p*) *for each p* ∈ *P, such that P covers all multiple-end phasing paths in 𝒞, and that both d*(*P, f*) *and* |*P*| *are as small as possible*.

We note that above informal description is similar to the formulation we used in Scallop [7], as the principles of parsimony and minimizing deviation apply to both cases. More importantly, both formulations aim to fully make use the long-range information in the data (for multiple-end RNA-seq data such information is abstracted as multiple-end phasing paths, and for short-reads data such information is abstracted as single-end phasing paths), and this is achieved by enforcing the long-range structures (either multiple-end phasing paths or single-end phasing paths) to appear in the assembled transcripts.

### 3.3 Algorithmic framework

We propose a heuristic for Problem 1. This heuristic consists of three major steps, described below.

**Step 1: bridging multiple-end phasing paths into single-end phasing paths**. Let *C* = {*l*_1_, *l*_2_, …, *l*_*n*_} ∈ *𝒞* be a multiple-end phasing path. We sort all paths {*l*_1_, *l*_2_, …, *l*_*n*_} in *C* in lexicographical order w.r.t. a topological ordering of all vertices of the splice graph (recall that the splice graph is a directed acyclic graph). See Figure 6. We still write *C* = (*l*_1_, *l*_2_, …, *l*_*n*_) after sorting all the paths lexicographically. All paths in *C* can be regarded as being sampled from a single (unknown) *s*-*t* path of the splice graph, representing the true transcript from which the reads used to construct *C* are generated. We propose to “bridge” all paths in *C* as a single-end phasing path, denoted as *h*(*C*), in the splice graph. This task amounts to filling the gaps (if any) between adjacent paths in *C* (see Figure 6). Specifically, if *l*_*i*_ and *l*_*i*+1_ overlap, i.e., there exists an suffix of *l*_*i*_ that is also a prefix of *l*_*i*+1_, then *l*_*i*_ and *l*_*i*+1_ can be naturally concatenated into a single path; otherwise, we will need to fill the gap by inferring a path connecting *l*_*i*_ to *l*_*i*+1_ in the splice graph (see below). After all *n* − 1 pairs of adjacent paths in *C*, i.e., (*l*_1_, *l*_2_), (*l*_2_, *l*_3_), …, (*l*_*n*−1_, *l*_*n*_), are bridged, we can then connect them together as a single-end phasing path, i.e., *h*(*C*). The algorithm to infer the true path that connects non-overlapping *l*_*i*_ and *l*_*i*+1_ is one major innovation in Scallop2, separately described in Section 3.4.

**Figure 6:**
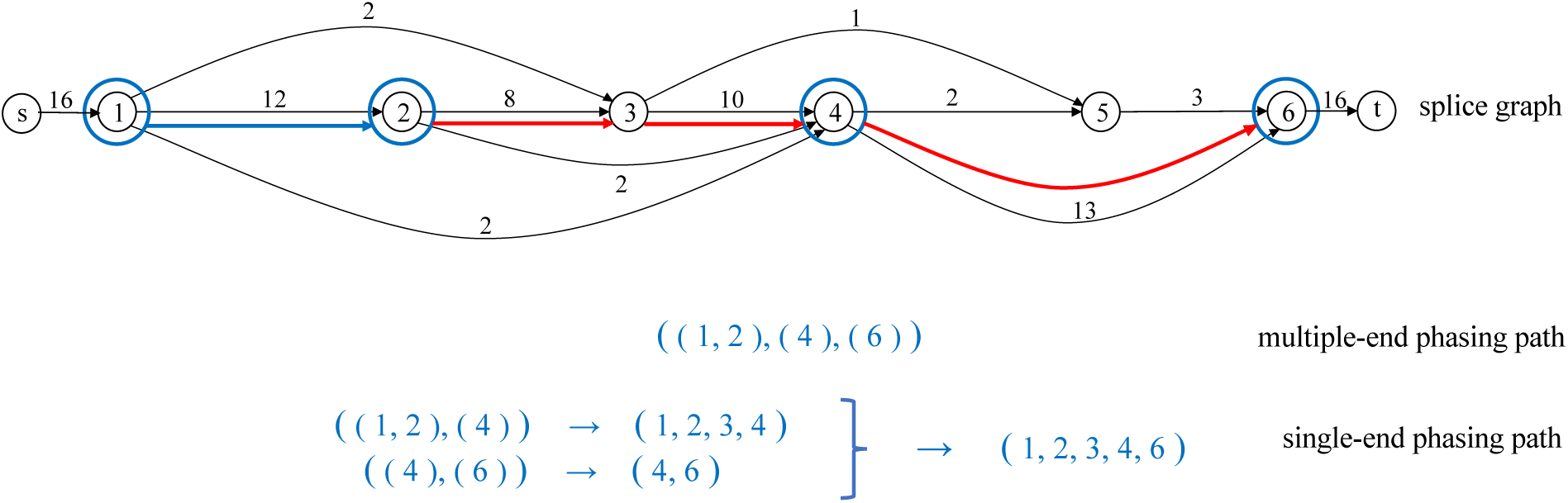
Illustration of bridging a multiple-end phasing path *C* = (*l*_1_ = (1, 2), *l*_2_ = (4), *l*_3_ = (6)) into a single-end phasing path *h*(*C*) = (1, 2, 3, 4, 6). Inferred bridging paths for (*l*_1_, *l*_2_) and (*l*_2_, *l*_3_) are marked red.

**Step 2: determining false starting/ending vertices**. We observe that failing to bridge a multiple-end phasing path is mainly because the underlying splice graph is incomplete, such as certain junctions are missing due to low coverage or alignment errors. We design a new algorithm to determine false starting and ending vertices in the splice graph, separately described in Section 3.5. This algorithm constitutes another algorithmic innovation of Scallop2.

**Step 3: decomposing splice graph**. Let *H* := {*h*(*C*) | *C* ∈ *𝒞*} be the set of bridged single-end phasing paths returned by Step 1. Let *G*′ be the refined splice graph returned by Step 2. Notice that now the objective of preserving all multiple-end phasing path in *𝒞* becomes a simpler constraint of preserving all single-end phasing paths in *H*. In our previous work Scallop [7], we have designed an efficient heuristic that solves *exactly* the same question of decomposing the splice graph in the presence of (single-end) phasing paths. Here, we directly use this algorithm to obtain the set of *s*-*t* paths *P* by piping *G*′ and *H* into this algorithm.

### 3.4 Bridging algorithm

Let *l* = (*a*_1_, *a*_2_, …, *a*_*m*_), *l*′ = (*b*_1_, *b*_2_, …, *b*_*n*_) be any two consecutive, non-overlapping paths in a multiple-end phasing path. The problem of bridging *l*_*i*_ and *l*_*i*+1_ amounts to find a path in *G* from *a*_*m*_ to *b*_1_. Such path, called *bridging path*, infers the missing portion between *l*_*i*_ and *l*_*i*+1_. We note that there might be multiple bridging paths due to alternative splicing and sequencing/alignment errors and we aim to infer the correct one. Below we first formulate the task of inferring the true bridging path as a new optimization problem and then design an efficient algorithm.

#### Formulation

We explore what’s a good formulation to find the true bridging path. The main signal we have is the coverage information, and intuitively a bridging path supported by most reads are most likely the true path. We determined that, a formulation that seeks a bridging path that maximizes *bottleneck* weight, is suitable for this bridging problem. Below we formally describe this formulation.

We define a full ordering of all bridging paths. Let *q*_1_ and *q*_2_ be two arbitrary paths from *a*_*m*_ to *b*_1_ in *G*. Let 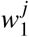 (resp. 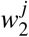) be the *j*th smallest weight in path *q*_1_ (resp. *q*_2_). We say *q*_1_ is *more reliable* than *q*_2_, if there exists an integer *k* such that 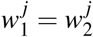 for all 1 ≤ *j < k*, and 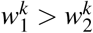. In other words, each bridging path is represented as the *sorted* list of its weights in ascending order, and all bridging paths are then (implicitly) sorted in lexicographical order. We formulate the problem of bridging as to find the most reliable path. Intuitively, we seek a path *q* from *a*_*m*_ to *b*_1_ in *G* such that the smallest weight in it is maximized; in case there are multiple paths with maximized smallest weight, among them we further seek the one whose second smallest weight is maximized, and so on.

We believe this formulation is appropriate for RNA-seq bridging. First, by maximizing bottleneck weight, the bridging paths are supported strongest, and hence are more likely to be the true bridging path. Second, through this formulation, false paths which are usually due to sequencing/alignment errors and therefore exhibit low abundances, can be automatically and efficiently excluded.

We note that this formulation satisfies the *optimal substructure* property: if *a*_*m*_ → *u*_1_ → *u*_2_ → … → *u*_*l*_ → *b*_1_ is the most reliable path from *a*_*m*_ to *b*_1_, then *a*_*m*_ → *u*_1_ → *u*_2_ → … → *u*_*l*_ is the most reliable path from *a*_*m*_ to *u*_*l*_. This allows us to design a dynamic programming algorithm to find the most reliable bridging path.

#### Dynamic programming algorithm

Let (*v*_1_, *v*_2_, …, *v*_|*V*|_) be a topological sorting of all vertices of the splice graph *G* (this is possible because splice graph is a directed acyclic graph). Given a particular *v*_*i*_, we can use a single run to find the most reliable paths from *v*_*i*_ to *v* _*j*_ for every *j > i*. To compute the optimal path for *v* _*j*_, we examine all vertices *v*_*k*_ that directly connects to *v* _*j*_, and compare all paths stored in these vertices (each *v*_*k*_ already stores the most reliable paths from *v*_*i*_ to *v*_*k*_ at this time point). The optimal one will be kept and after concatenating *v* _*j*_ they become the most reliable paths from *v*_*i*_ to *v* _*j*_. We run this subroutine for all *v*_*i*_, 1 ≤ *i* ≤ |*V* |, which gives the most reliable paths for all pairs of vertices in *G*. The overall running time of this algorithm is *O*(|*V* |2 · |*E*|). To speed up, instead of maintaining the full list of the edge abundances for each path, whose length is *O*(|*V* |), we only store the smallest *M* edge abundances (*M* is a parameter with default value of 5). This gives an improved running time of *O*(*M* · |*V* | · |*E*|). Although optimality may not be guaranteed, experimental studies show that this heuristic rarely affects the overall accuracy.

#### Determining whether or not to accept the most reliable path

In case of paired-end RNA-seq data, the distribution of fragment length provides another source of information to decide if the inferred most reliable path is correct. Scallop2 will estimate the distribution of fragment length by sampling reads that can be trivially bridged (i.e., they overlap in the splice graph). Once we determine the most reliable path connecting *l*_1_ and *l*_2_ in a (paired-end) phasing path *C* = (*l*_1_, *l*_2_), we can also calculate the fragment length, as the alignment of the entire fragment is fixed. If the resulting fragment length falls in a *reasonable range*, by default defined as the 0.5-percentile to 99.8-percentile of the estimated distribution, then the inferred path will be accepted; otherwise we mark that *C* is failed to bridge.

### 3.5 Determining false starting/ending vertices

The constructed splice graph is usually erroneous due to missing junctions, sequencing and alignment errors. We observe that erroneous starting and ending vertices (i.e., those connected to the source and sink vertices, respectively) can be identified through reads that are failed to bridge. Figure 7 gives an example: the two blue reads and the second and the third red reads cannot be bridged, as there is no edge connecting vertices 2 and 3; these suggest that vertices 2 and 3 are false starting and ending vertices (there is likely a missing junction between vertices 2 and 3 in this example).

**Figure 7:**
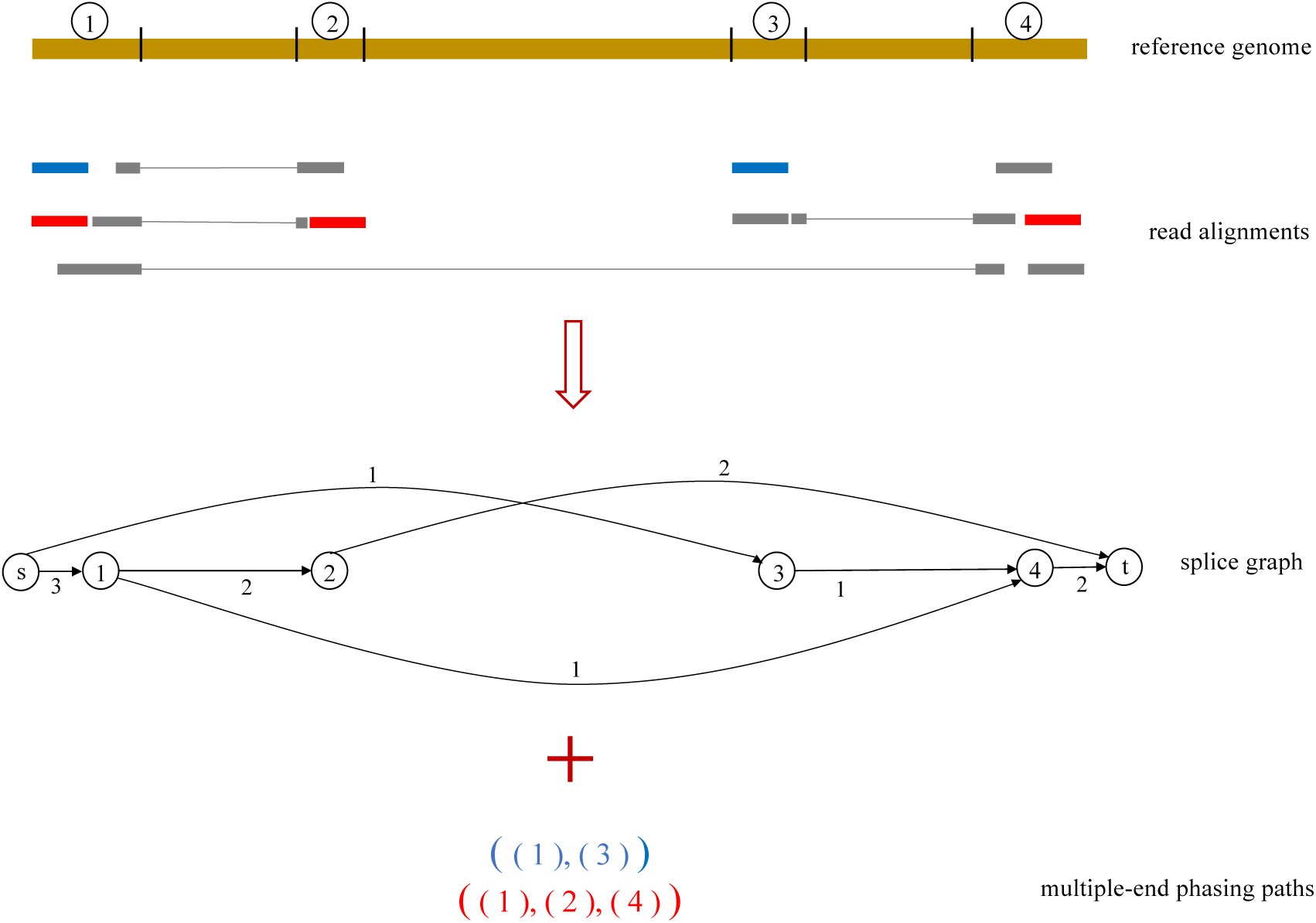
Illustration of identifying false starting/ending vertices. From the given alignments 4 partial-exons (numbered 1–4) are identified. Vertex 2 (resp. 3) is classified as ending (resp. starting) vertex as there is no departing (resp. entering) junction. An pseudo-edge (2, 3) will be added to the splice graph for bridging. The bridging of blue reads and red reads (the second and the third ones) will cross this pseudo-edge, giving a pseudo-score of 2 for both vertices.

We design an algorithm that implements above observation. Let *u* and *v* be two consecutive vertices in the splice graph (i.e., there is no any other vertex between *u* and *v*). If *u* is an ending vertex (i.e., (*u,t*) ∈ *E*) and *v* is a starting vertex (i.e., (*s, u*) ∈ *E*), then we will add a *pseudo-edge* (*u, v*) with weight of 0.5 to the splice graph. (In Figure 7, edge (2, 3) will be added as psuedo-edge.) The expanded splice graph (with pseudo-edges added) will be actually used for bridging all consecutive ends by the algorithm described in Section 3.4. Notice that any non-pseudo-edge will have a weight at least 1; therefore, if the bottleneck-weight of the most reliable path *p* of a pair of ends (*l, l*′) equals to 0.5, then we know that *p* must cross at least one pseudo-edge and that (*l, l*′) cannot be bridged with non-pseudo-edges only. In this case, we will examine all pseudo-edges (*u, v*) in *p*, and assign a *pseudo-score* of value 1 to *u* and to *v*. We run this algorithm for all read groups, and the pseudo-score will be accumulated. (For example, in Figure 7, both vertices 2 and 3 will have a pseudo-score of 2; the blue and the red reads contribute 1 each.) Intuitively, the larger pseudo-score is, the more likely the vertex is a false starting/ending vertex. Let *x*(*v*) be the pseudo-score of vertex *v* and let *w*(*v*) be the average coverage of *v*. We calculate *z*(*v*) := log(*w*(*v*)+1)−log(*x*(*v*)+1) and report *v* is a false vertex only if *x*(*v*) ≥ 1 and *z*(*v*) is less than a threshold (by default 1.5). We do this to ensure that, starting/ending vertices with large coverage will be determined as false only if there exists a relatively significant number of supporting reads.

The determined false starting and ending vertices will be kept in the splice graph for decomposition (Step 3; Section 3.3). A resulting *s*-*t* path that contains any false vertex will be classified as “transcript fragment” (as opposed to “full-length transcript”). We keep these transcript fragments as they explain the reads in the false vertices (it’s that we have evidence to determine they are not full-length transcripts). In our implementation full-length transcripts and transcript fragments will be written to separate files. The accuracy of assemblies reported in Section 2 are for the full-length transcripts.

## Acknowledgments

This work is partly supported by the US National Science Foundation (DBI-2019797 to M.S.), by the US National Institutes of Health (R01HG011065 to M.S.), and by the Charles K. Etner Career Development Professorship awarded to M.S. by The Pennsylvania State University. Initial algorithmic exploration of Scallop2 advancements were conducted with Carl Kingsford and were supported by the Gordon and Betty Moore Foundation (GMBF 4554 to C.K.) and the US National Institutes of Health (R01GM122935 to C.K.).

## Appendix

We were not able to compare with scRNAss in our experiments. The raw RNA-seq data is demultiplexing and it requires to be processed by zUMIs to distinguishes cell barcodes and molecular barcodes from sequencing reads, which automatically generates STAR-aligned bam files. However, for the current version of scRNAss, it only identifies the fastq/fasta files as inputs. We were not compare with RNA-Bloom in our experiments, because in RNA-Bloom’s reference-guided mode, k-mer pairs from a transcriptome reference are included in addition to those derived from reads and fragments’, which means that RNA-Bloom uses reference transcriptome instead of alignments to the reference genome.

